# Heart generation via blastocyst complementation in Mesp1/2-deficient mice

**DOI:** 10.1101/2023.11.08.565924

**Authors:** Yuto Abe, Taketaro Sadahiro, Ryo Fujita, Yu Yamada, Tatsuya Akiyama, Koji Nakano, Seiichiro Honda, Yoko Tanimoto, Hayate Suzuki, Seiya Mizuno, Satoru Takahashi, Tomoyuki Yamaguchi, Hideki Masaki, Kosuke Hattori, Tomoji Mashimo, Masaki Ieda

## Abstract

Heart transplantation is the only curative option available for patients with advanced heart failure. However, donor organ shortage and graft rejection remain critical challenges in heart transplantation. Blastocyst complementation using pluripotent stem cells (PSCs) can be used to generate organs such as the pancreas and kidneys in animal models with abnormalities in essential developmental genes. Nonetheless, whether functional adult hearts can be generated with blastocyst complementation remains unclear, and it is unknown if parenchyma, blood vessels, and stroma can be generated concomitantly from PSCs using blastocyst complementation to avoid graft rejection. Here, we show the generation of functional adult hearts in acardiac Mesp1 and Mesp2 double-knockout (Mesp1/2-DKO) mice via blastocyst complementation using mouse PSCs. Our result shows that the generated hearts were structurally and functionally normal and restored embryonic lethality in Mesp1/2-DKO mice. All four cardiovascular lineages, including cardiomyocytes, vascular endothelial cells, smooth muscle cells, and cardiac fibroblasts, were virtually entirely derived from exogenous PSCs in the myocardium. Exogenous rat PSCs also generated rat-derived xenogeneic hearts in Mesp1/2-DKO mice via interspecies blastocyst complementation. Thus, blastocyst complementation is a viable technique for generating hearts derived from PSCs and may represent significant progress toward generating rejection-free hearts.

## Introduction

Heart failure (HF) is a life-threatening condition worldwide. The prognosis has improved with advancements in pharmacological and non-pharmacological therapies; nonetheless, heart transplantation is the only curative treatment for severe HF^1^. However, the quantity of donor hearts for patients with HF is insufficient, and novel treatments are eagerly anticipated^2^. Cell-based therapy using pluripotent stem cell (PSC)-derived cardiomyocytes (CMs) can be an appealing option; nevertheless, issues, such as the immaturity of *in vitro* generated CMs, risk of tumorigenesis due to contamination with undifferentiated cells, difficulties in transplanted cell engraftment, and moderate effects on functional recovery, remain^3,4^.

Blastocyst complementation is a potential method for generating the desired organs^5^. In intraspecies and interspecies chimeras, injection of PSCs into blastocysts lacking Pdx1, a transcription factor critical for pancreatic development, generates a functional pancreas composed of PSC-derived endocrine and exocrine cells ^6^. In addition, transplantation of mouse PSC-derived pancreatic islets generated in Pdx1-deficient rats successfully restored blood glucose levels in diabetic mice for more than one year without the need for continuous immunosuppression^7^. PSC-derived kidneys and lungs were also generated in anephric Sall1- and apneumic Fgf10-deficient animals, respectively^8,9^. However, it is unknown if using blastocyst complementation can yield functional adult hearts.

The heart consists of CMs and a large number of non-CMs, including vascular endothelial cells (ECs), smooth muscle cells (SMCs), and cardiac fibroblasts (CFs)^10^. Non-CMs accurately coordinate with CMs to maintain cardiac circulation and function^11^. Moreover, non-CMs are responsible for rejection following organ transplantation; a major histocompatibility complex (MHC) mismatch in ECs can result in hyperacute rejection in transplanted hearts^12–14^. Therefore, it is critical to generate hearts consisting of CMs and non-CMs produced solely from exogenous PSCs using blastocyst complementation. However, the vascular cells and mesenchyme in the pancreas and kidneys formed using blastocyst complementation are mixes of host- and donor-derived cells ^6,7,9^. It remains unclear whether blastocyst complementation can be used to generate parenchyma (CMs) and stroma (non-CMs) from PSCs concomitantly.

Mesp1 and Mesp2 are adjacent transcription factors that regulate mesodermal specification and cardiovascular differentiation in a redundant manner^15–18^. Mesp1 and Mesp2 double-knockout (Mesp1/2-DKO) mice die at embryonic day (E) nine because of defects in mesoderm formation and its derivatives, including hearts^17–19^. MESP1 deficiency in human PSCs inhibits epithelial-mesenchymal transition and hinders cardiac differentiation, highlighting the importance and conserved functions of Mesp1 and 2 in cardiogenesis ^20^. Using mouse and rat PSCs, we investigated whether blastocyst complementation could be used to generate hearts in Mesp1/2-DKO mice.

## Results

### Mesp1/2-DKO mice provide organ niches for heart generation via blastocyst complementation

By exploiting the developmental organ niche, blastocyst complementation can be used to generate organs from PSCs. Mesp1 is a key regulator of cardiovascular differentiation that is expressed transiently in the cardiac mesoderm^16,19,20^. We analyzed Mesp1-Cre/R26-tdTomato mice to determine the cell lineages formed from Mesp1^+^ cells. Notably, the entire myocardium and all four cardiovascular lineages (CMs, ECs, SMCs, and CFs) expressed tdTomato, suggesting that they were derived from the Mesp1^+^ mesoderm (Extended Data Fig. 1a, b). Previous studies have revealed that Mesp1 and 2 function redundantly in cardiovascular development, and Mesp1/2-DKO mice exhibit mesodermal abnormalities, but not Mesp1-KO or Mesp2-KO mice^17,18^. Consistent with the findings above, we found that Mesp1/2-DKO (Mesp1^−/−^/Mesp2^−/−^, Mesp^−/−^) mice were embryonically lethal at E9.0 with defects in heart formation and posterior trunk anatomy (Fig. 1a, b). qRT-PCR analysis revealed that cardiac genes *Myh6* and *Tnnt2* expression were eliminated in Mesp^−/−^ embryos (Fig. 1c).

**Fig. 1.**
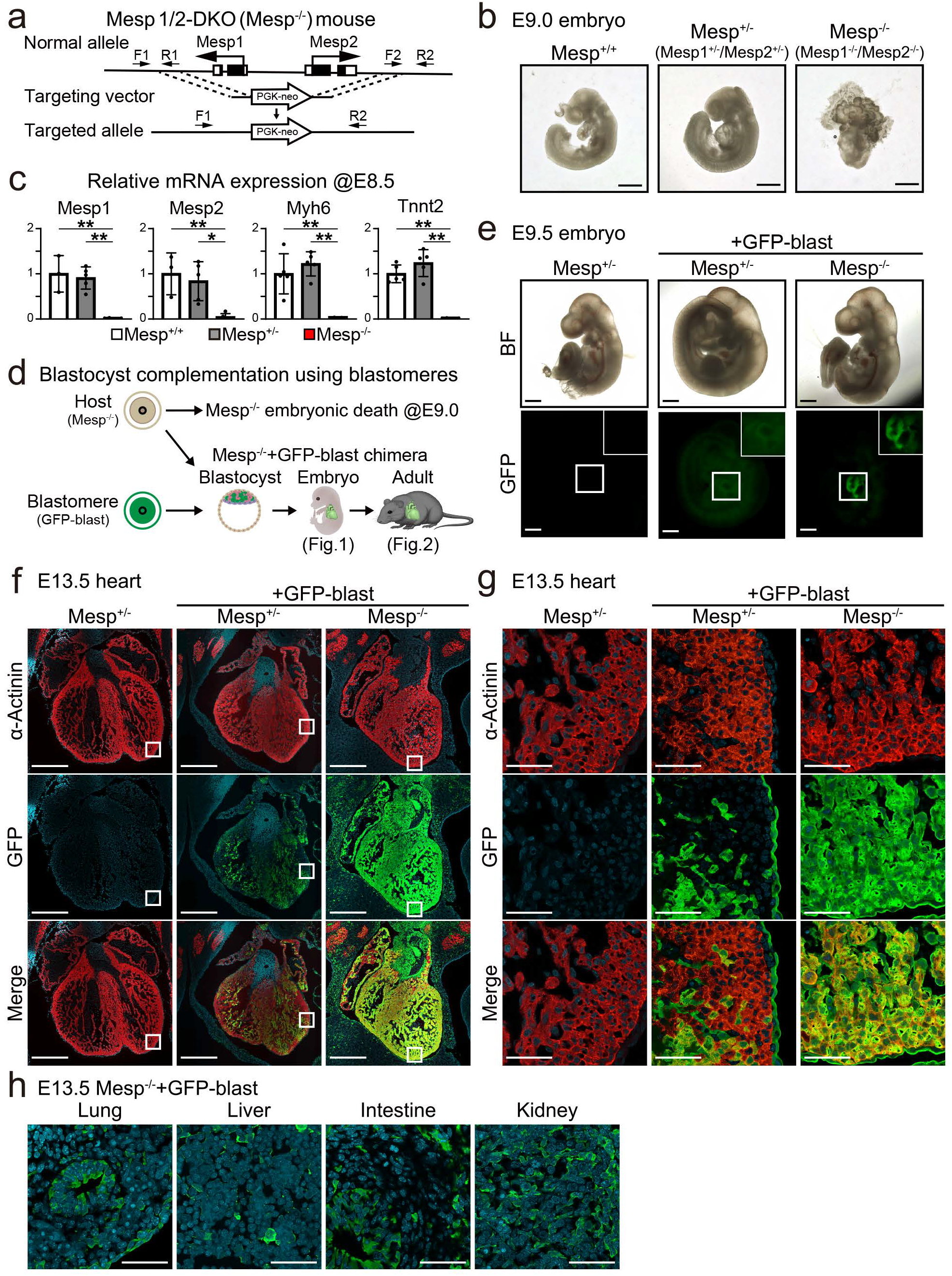
Blastocyst complementation in Mesp1/2-DKO mice using blastomeres. **a,** Scheme of Mesp1 and Mesp2 double knockout mouse (Mesp1/2-DKO: Mesp1^−/−^/Mesp2^−/−^, Mesp^−/−^). Mesp^−/−^ mice were obtained by crossing Mesp1/2 heterozygous KO mice (Mesp1^+/−^/Mesp2^+/−^, Mesp^+/−^). F1, F2, R1, and R2 indicate the locations of the PCR primers used for genotyping. **b,** Representative whole-mount images of Mesp^+/+^, Mesp^+/−^, and Mesp^−/−^ mice at embryonic day (E) 9.0. Scale bars, 500 μm. **c,** Relative mRNA expression of cardiac-related genes at E8.5. The graph represents mean ± SD of *Mesp1, Mesp2*, *Myh6*, and *Tnnt2* mRNA expression; each n=3~5 embryos per group. One-way ANOVA test with Tukey’s post-hoc test was used for statistical analysis. \**P*<0.05, \*\**P*<0.01 **d,** Schematic of the blastocyst complementation strategy using GFP-labeled blastomere (GFP-blast) in Mesp^−/−^ mice. **e,** Brightfield and fluorescence images of embryos at E9.5. High-magnification images of the heart are shown in the insets. Scale bars, 500 μm. **f, g,** Immunohistochemistry for α-actinin, GFP, and DAPI in E13.5 embryonic hearts (**f**). High-magnification images of the white squares are shown in (**g**). Scale bars: (**f**) 500 μm, (**g**) 100 μm. **h,** Immunofluorescence images of GFP and DAPI in the lungs, liver, intestine, and kidneys of Mesp^−/−^ mice complemented with GFP^+^-blastomeres. Scale bars, 100 μm.

Using totipotent blastomeres, we then performed blastocyst complementation of Mesp^−/−^ embryos (Fig. 1d). Mesp^−/−^ mice were obtained by intercrossing Mesp1/2 heterozygous KO mice (Mesp1^+/−^/Mesp2^+/−^, Mesp^+/−^) that were viable and had a normal cardiac function. Mutant embryos were aggregated with a green fluorescence protein (GFP)-labeled wild-type blastomeres (GFP-blast) and transferred to pseudopregnant females at the blastocyst stage (Fig. 1d). Each genotype was validated by PCR in chimeras, using genomic DNA extracted from fluorescence-activated cell sorting (FACS)-isolated GFP-negative host fibroblasts (Extended Data Fig. 2a, b, 3a). At E9.5, 18 chimeric mice were obtained, of which two embryos were Mesp^−/−^ + GFP-blasts. In contrast to Mesp^−/−^ embryos, Mesp^−/−^ + GFP-blast mice grew normally, similar to the control embryos (Mesp^+/−^ with or without GFP-blasts). Importantly, Mesp^−/−^ + GFP-blast hearts strongly expressed GFP, suggesting that they were generated from GFP blastomeres (Fig. 1e). At E13.5, immunohistochemistry (IHC) revealed that the myocardium in Mesp^+/−^ + GFP-blast consisted of a mixture of GFP^+^ and GFP^−^ cells (the ratio of GFP^+^ cells: 50.6% ± 7.6%, mean ± SD), whereas that in Mesp^−/−^ + GFP-blast was predominantly composed of GFP^+^ cells (91.7% ± 3.9%) (Fig. 1f, g). Further, other organs in the Mesp^−/−^ + GFP-blast, such as the lungs, liver, intestine, and kidneys, had GFP^+^ and GFP^−^ cell composites (GFP^+^ cells in each organ: 53.4% ± 11.5%, 37.1% ± 6.2%, 52.9% ± 10.2%, and 59.6% ± 12.7%) (Fig. 1h). These findings suggest that cardiac defects in Mesp1/2-DKO mice are cell autonomous. In contrast, other organ developmental defects are most likely non-cell autonomous. Thus, Mesp1/2-DKO mice provide an organ niche for heart generation via blastocyst complementation.

### Functional adult heart generation in Mesp1/2-DKO mice via blastocyst complementation using blastomeres

Next, we analyzed Mesp^−/−^ blastocyst complementation using GFP-blasts in 8-week-old mice. Overall, 100 blastocysts were transplanted into pseudopregnant mouse uteri, and 35 adult mice were obtained. Twenty-five of these were identified as chimeras based on GFP expression. Genotyping revealed that five of the 25 chimeras (20%) were Mesp^−/−^ + GFP-blast mice, suggesting that blastocyst complementation may rescue Mesp^−/−^ mice into adulthood (Extended Data Fig. 3a). Their body sizes were similar; however, all Mesp^−/−^ + GFP-blast mice had short and kinky tails reminiscent of the phenotype of Mesp2^−/−^ mice (Fig. 2a, c)^21^. Furthermore, IHC revealed that hearts in Mesp^+/−^ + GFP-blast mice were a mix of host- and donor-derived cells, similar to whole-body chimerism (Fig 2b, Extended Data Fig. 3b). In contrast, hearts in Mesp^−/−^ + GFP-blast mice were almost exclusively composed of GFP^+^ cells (Fig. 2d). Co-immunostaining of GFP and multiple markers indicated that CMs and non-CMs in Mesp^+/−^ + GFP-blast were a composite of host and donor derivatives (Fig. 2e). In contrast, in Mesp^−/−^ + GFP-blast hearts, CMs, ECs, SMCs, and CFs were virtually exclusively generated from GFP^+^ blastomeres (Fig. 2f). In Mesp^−/−^ + GFP-blast mice, other organs were composites of GFP^+^ and GFP^−^ cells, suggesting that complementation is exclusive to heart formation (Fig. 2g). Mesp^−/−^ + GFP blast hearts were morphologically and histologically normal (Extended Data Fig. 3c-e). To determine cardiac function, we performed echocardiography and treadmill tests on control (Mesp^+/+^ and Mesp^+/−^ + GFP-blast) and Mesp^−/−^ + GFP-blast mice. The left ventricular ejection fraction was comparable across the three groups, and exercise capability was maintained in Mesp^−/−^ + GFP-blast mice (Fig. 2h, i, Extended Data Fig. 3f, g). These findings suggested that blastocyst complementation in Mesp^−/−^ mice generates functioning adult hearts that are almost entirely generated from exogenous blastomeres.

**Fig. 2.**
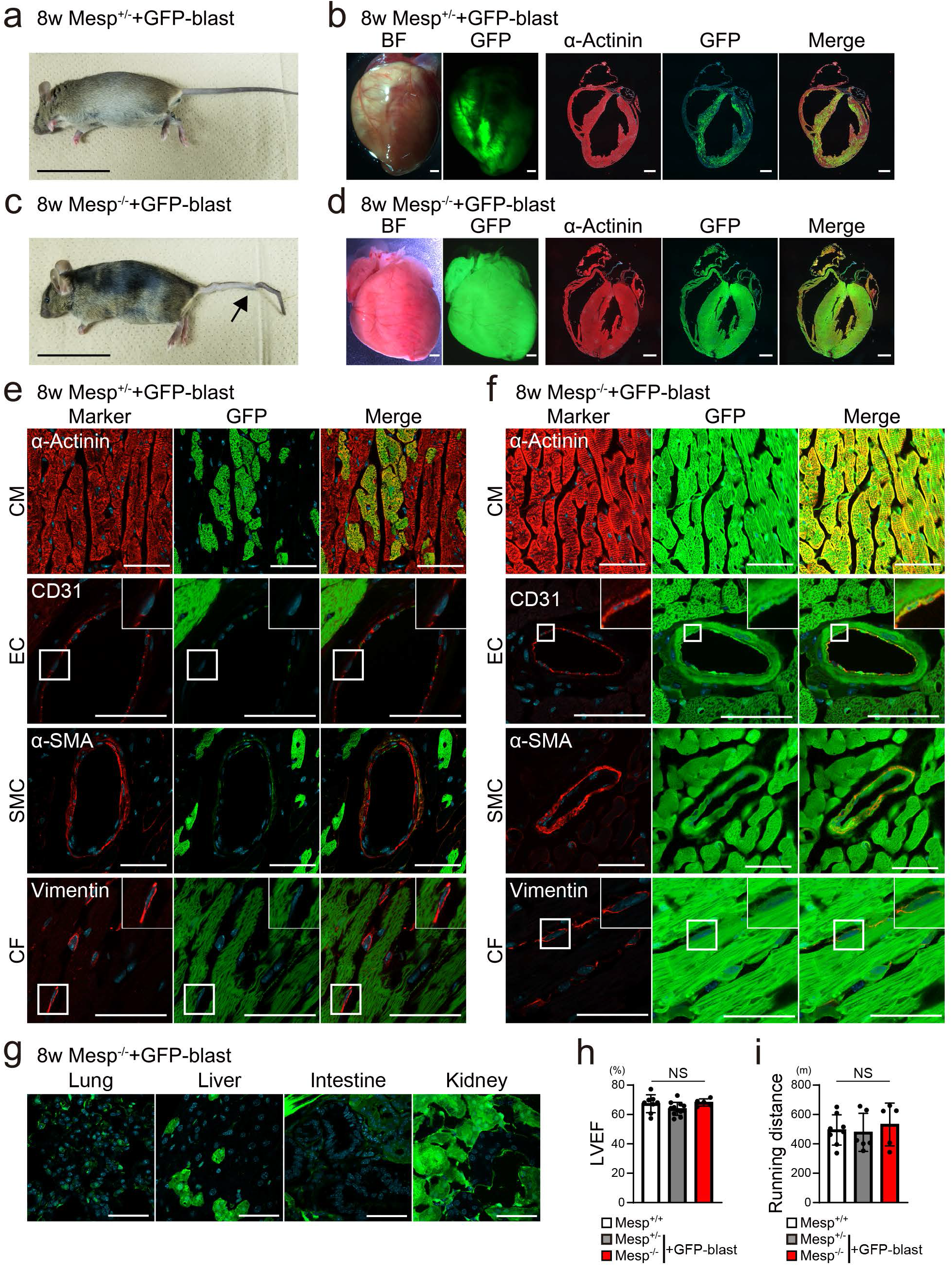
Generation of functional adult hearts by blastocyst complementation using blastomeres. **a, c,** Representative images of adult Mesp^+/−^ + GFP-blast (**a**) and Mesp^−/−^ + GFP-blast (**c**) chimeric mice at 8 weeks of age. The arrow indicates the kinked tail. Scale bars, 5 cm. **b, d,** Whole-mount brightfield and GFP fluorescence images, and sections of Mesp^+/−^ + GFP- blast (**b**) and Mesp^−/−^ + GFP-blast (**d**) chimeric hearts. Heart sections were immunostained with α-actinin, GFP, and DAPI. Scale bars, 1 mm. **e, f,** Immunohistochemistry for the expression of cardiovascular cell markers, GFP and DAPI, in Mesp^+/−^ + GFP-blast (**e**) and Mesp^−/−^ + GFP-blast chimeric hearts (**f**). Antibodies against actinin, a marker of cardiomyocytes (CM), CD31 for endothelial cells (EC), αSMA for smooth muscle cells (SMC), and vimentin for cardiac fibroblasts (CF) were used. High-magnification images are shown in the insets (EC and CF). **g,** Immunofluorescence images of GFP and DAPI in the lungs, liver, intestine, and kidneys of adult Mesp^−/−^ + GFP blast chimeras. Scale bars, 100 μm. **h,** Left ventricular ejection fraction (LVEF) was measured using echocardiography. Graph represents mean ± SD of n=8 (Mesp^+/+^), n=12 (Mesp^+/−^ + GFP-blast), and n=5 (Mesp^−/−^ + GFP-blast) mice per group. One-way ANOVA test was used for statistical analysis. NS: statistically non-significant. **i,** Running distance during the treadmill fatigue test. Graph represents mean ± SD of n=8 (Mesp^+/+^), n=6 (Mesp^+/−^ + GFP-blast), and n=5 (Mesp^−/−^ + GFP-blast) mice per group. One-way ANOVA test with Tukey’s post-hoc test was used for statistical analysis. NS: statistically non-significant

### Blastocyst complementation in Mesp1/2-DKO mice using mouse embryonic stem cells

The generation of functional adult hearts via blastocyst complementation using blastomeres is remarkable; however, ethical concerns may prevent its clinical application. We used GFP-labeled wild-type mouse embryonic stem cells (GFP-mESCs) to execute Mesp^−/−^ blastocyst complementation. GFP-mESCs were injected into blastocysts derived from intercrossing heterozygous Mesp^+/−^ mice (Fig. 3a, Extended Data Fig. 4a). We obtained two Mesp^−/−^ + GFP-mESCs with high GFP expression in the heart at E9.5 (Extended Data Fig. 4a, b). The transfer of 172 supplemented blastocysts to pseudopregnant females resulted in 51 fetuses at E13.5, 34 of which were chimeric and 17 non-chimeric fetuses. Five of the 34 chimeras were Mesp^−/−^ + GFP-mESCs with GFP-expressing hearts (Fig. 3b, Extended Data Fig. 4a). Mesp^−/−^ + GFP-mESCs grew normally, as did the controls (Mesp^+/−^ with or without GFP-mESCs) (Fig. 3b). IHC revealed that the myocardium of Mesp^−/−^ + GFP-mESC mice was almost exclusively consisted of GFP^+^ cells, whereas that of Mesp^+/−^ + GFP-mESCs was a mix of GFP^+^ and GFP^−^ cells (Fig. 3c). The other organs in the Mesp^−/−^ + GFP-mESCs were composed of host- and donor-derived cells (Extended Data Fig. 4c).

**Fig. 3.**
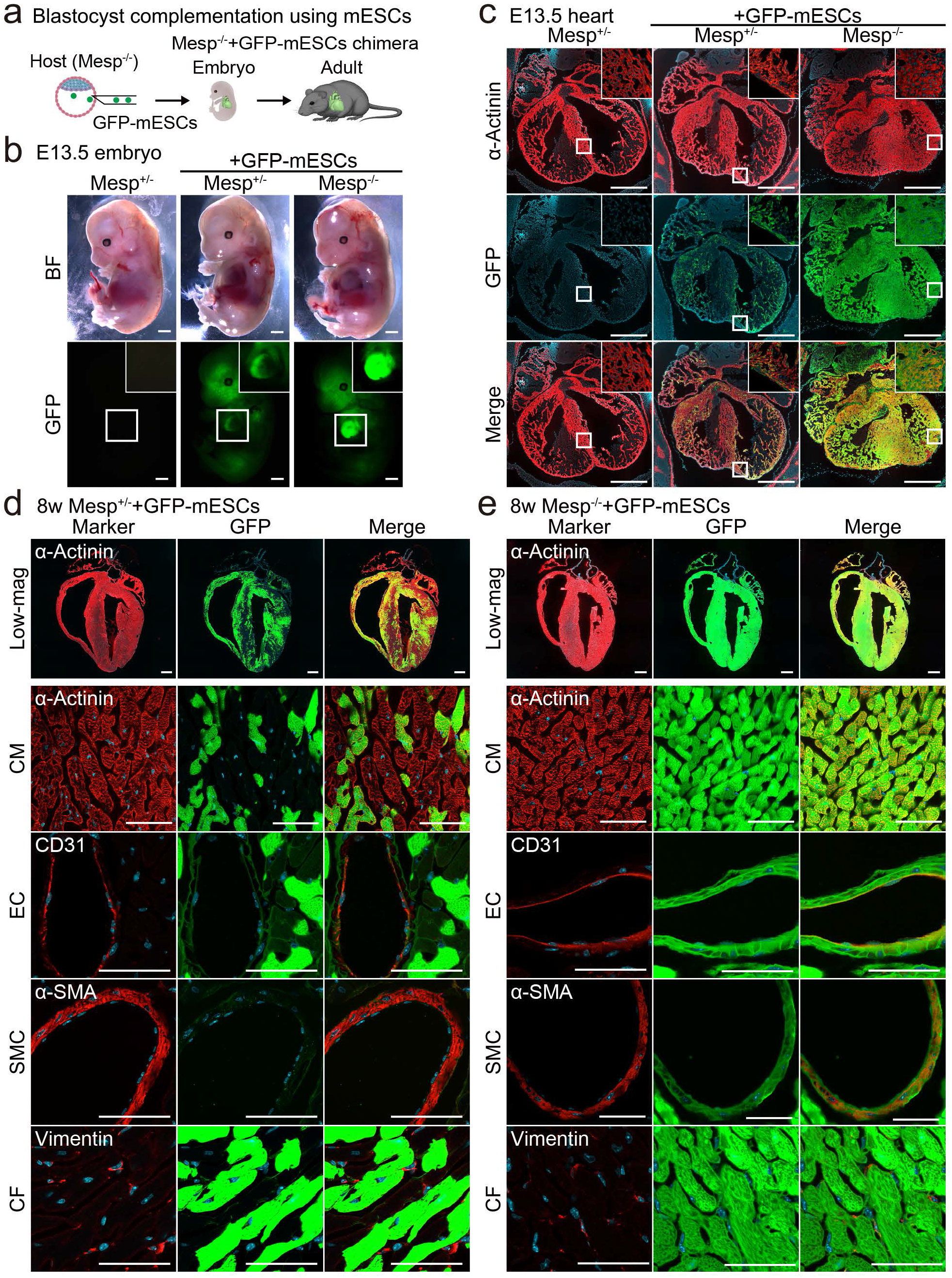
Blastocyst complementation in Mesp1/2-DKO mice using mouse embryonic stem cells. **a,** Schematic of the blastocyst complementation strategy in Mesp^−/−^ mice using GFP-labeled mouse embryonic stem cells (GFP-mESCs). **b,** Brightfield and fluorescence images of embryos at E13.5. High-magnification images of the heart are shown in the inset. Scale bars, 1 mm. **c,** Immunohistochemistry (IHC) for α-actinin, GFP, and DAPI in embryonic hearts at E13.5. High-magnification images are shown in the insets. Scale bars, 500 μm. **d, e,** Upper-panels; IHC for α-actinin, GFP and DAPI in Mesp^+/−^ + GFP-mESCs (**d**) and Mesp^−/−^ + GFP-mESCs mouse hearts (**e**). Whole heart images are shown in a low-magnification view (Low-mag). Scale bars, 1 mm. Lower panels; IHC for the expression of cardiovascular cells (CM, EC, SMC, and CF), GFP, and DAPI in Mesp^+/−^ + GFP-mESCs (**d**) and Mesp^−/−^ + GFP-mESCs mouse hearts (**e**). Scale bars, 100 μm. CM, cardiomyocyte; EC, endothelial cell; SMC, smooth muscle cell; CF, cardiac fibroblast

We then analyzed Mesp^−/−^ blastocyst complementation using GFP-mESCs in 8-week-old adult mice. Overall, 625 blastocysts were transplanted into pseudopregnant mouse uteri, yielding 138 adult mice. Of these, 108 mice were identified as chimeras using GFP expression, with four of the 108 (3.7%) being Mesp^−/−^ + GFP-mESCs, suggesting that blastocyst complementation may be used to rescue Mesp^−/−^ mice into adulthood (Extended Data Fig. 4a). All of the Mesp^−/−^ + GFP-mESCs mice grew normally and the chimera mice had distinctive tails (Fig. 2c, Extended Data Fig. 5a, b). In addition, IHC revealed that Mesp^−/−^ + GFP-mESC hearts exhibited GFP evenly, whereas Mesp^+/−^ + GFP-mESC hearts expressed GFP patchily (Fig. 3d, e, Extended Data Fig. 5a, b). Quantitative analyses revealed that the cardiovascular cells, including CMs, ECs, SMCs, and CFs, were almost exclusively generated from GFP^+^ cells in Mesp^−/−^ + GFP-mESCs (GFP^+^ cells in each cell type: 95.0% ± 9.5%, 96.5% ± 3.8%, 94.7% ± 7.7%, and 91.6% ± 6.3%), whereas those of Mesp^+/−^ + GFP-mESCs were composites of GFP^+^ and GFP^−^ cells (GFP^+^ cells in each cell type: 42.4% ± 4.5%, 47.2% ± 9.7%, 46.2% ± 17.7%, and 39.1% ± 11.6%) (Fig. 3d, e). Other organs in both chimeras consisted of a mix of GFP^+^ and GFP^−^ cells (Extended Data Fig. 5c, d). Histological analyses of Mesp^−/−^ + GFP-mESC hearts revealed that they were structurally normal without tumorigenesis, hypertrophy, inflammation, or fibrosis (Extended Data Fig. 5e-g). Electrocardiography, echocardiography, and treadmill testing revealed that Mesp^−/−^ + GFP-mESC mice had normal cardiac function without arrhythmias (Extended Data Fig. 5h-j). Thus, exogenous mESCs supplemented the cardiac niche in Mesp^−/−^ mice, generating functional adult hearts almost exclusively derived from donor ESCs.

### Generation of a rat embryonic stem cell-derived xenogenic heart in Mesp1/2-DKO mice by interspecies blastocyst complementation

Next, GFP-labeled wild-type rat ESCs (GFP-rESCs) were injected into Mesp^−/−^ mouse blastocysts to create rat hearts (Fig. 4a). Further, because Mesp^−/−^ embryos died at E9.0, we analyzed the mice at E11.5 to determine the effect of interspecies blastocyst complementation. Overall, 153 supplemented blastocysts were transplanted to surrogate mothers, generating 50 embryos at E11.5. Nineteen of these were rat and mouse chimeras, with one (5%) being Mesp^−/−^ + GFP-rESCs, suggesting that interspecies blastocyst complementation may, albeit inefficiently, rescue Mesp^−/−^ mouse lethality (Extended Data Fig. 6). The body and heart sizes were similar between Mesp^−/−^ + GFP-rESCs and control embryos (Mesp^+/−^ with or without GFP-rESCs) (Fig. 4b, c). Mesp^−/−^ + GFP-rESC hearts strongly expressed GFP. IHC revealed that the heart, but not the somites or neural tubes, expressed GFP in Mesp^−/−^ + GFP-rESC mice (Fig. 4c). GFP was consistently expressed in the atria and ventricles of Mesp^−/−^ + GFP-rESCs homogenously, whereas Mesp^+/−^ + GFP-rESCs were composed of GFP^+^ and GFP^−^ cells (Fig. 4d, e). These findings revealed that by interspecies blastocyst complementation, rat ESCs could form a xenogeneic heart in Mesp^−/−^ mice.

**Fig 4.**
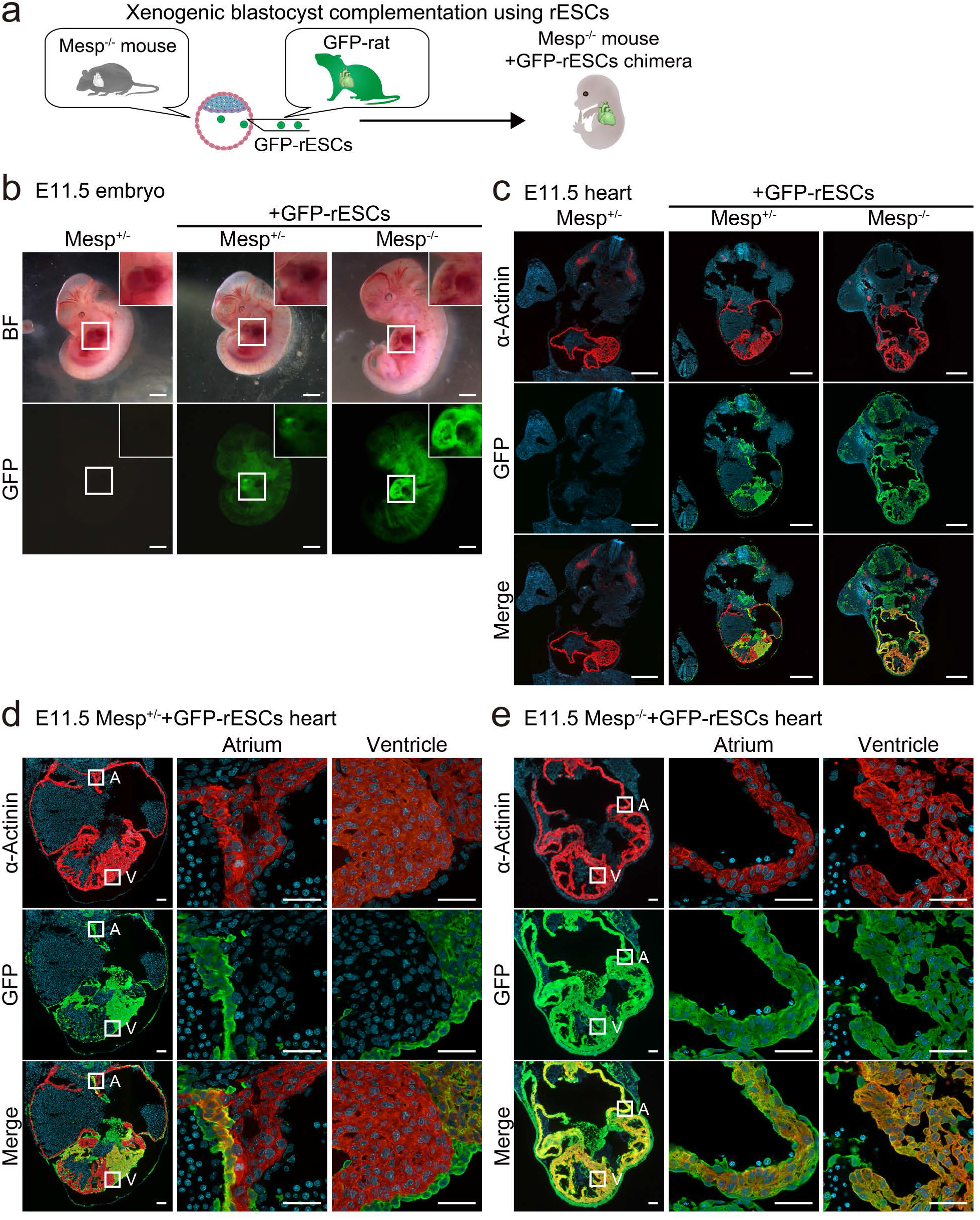
Generation of a rat ESC-derived xenogenic heart in Mesp1/2-DKO mouse by interspecies blastocyst complementation. **a,** Schematic representation of the interspecies blastocyst complementation strategy using GFP-labeled rat embryonic stem cells (GFP-rESCs). **b,** Bright-field and GFP fluorescence images of E11.5 embryos (Mesp^+/−^, Mesp^+/−^ + GFP-rESCs, and Mesp^−/−^ + GFP-rESCs). High-magnification images of the heart are shown in the inset. Scale bars, 1 mm. **c,** IHC for α-actinin, GFP, and DAPI in transverse sections of E11.5. Scale bars, 500 μm. GFP was mainly expressed in Mesp^−/−^ + GFP-rESC hearts. **d, e,** IHC for α-actinin, GFP, and DAPI in E11.5 chimeric hearts of Mesp^+/−^ + GFP-rESCs (**d**) and Mesp^−/−^ + GFP-rESCs (**e**). The left panels show low-magnification images. High-magnification images of the atria and ventricles are shown in the right panels. Scale bars, 100 μm. A, atrium; V, ventricle

## Discussion

Through blastocyst complementation using mouse PSCs, we were able to create functioning adult hearts in Mesp1/2-DKO mice. In the generated heart, CMs, ECs, SMCs, and CFs were almost exclusively derived from exogenous PSCs. Interspecies blastocyst complementation enables rat ESCs to create a xenogeneic heart in Mesp1/2-DKO mice. To our knowledge, this is the first study to reveal heart generation using mouse and rat PSCs via blastocyst complementation.

The heart comprises various cell types, such as CMs, ECs, SMCs, and CFs, which collectively work to maintain cardiac function. The complexity of heart organogenesis hinders *in vitro* generation of bone-fide hearts. This is the ultimate goal of regenerative medicine. Cell-based therapy has focused heavily on the direct *in vitro* differentiation of CMs from PSCs^22^; however, these CMs are functionally immature and arrhythmogenic, comparable to embryonic CMs^23,24^. Moreover, the possibility of tumor formation from PSC contamination and the poor survival of transplanted cells are significant challenges in clinical application^3,4^. In contrast, exploiting developmental niches in Mesp1/2-DKO mice, blastocyst complementation was used to create full adult hearts from PSCs. The CMs generated *in vivo* via blastocyst complementation were mature, similar to adult CMs, and the generated hearts functioned normally without tumorigenesis nor arrhythmias. A porcine heart genetically engineered to prevent xeno-rejection has recently been found to be transplantable into patients with end-stage heart failure^25^. Despite appropriate immunosuppressive therapy, treatment was withdrawn 60 d after transplantation because of xenograft failure. A xenotransplantation is an attractive approach for generating donor hearts; however, further improvements are needed^26^.

Blastocyst complementation generates the pancreas and kidneys, which consist of host- and donor-derived blood vessels and stroma^6,7,9^. It is critical to generate organs that are entirely derived from donor PSCs to avoid immunological rejection. Nkx2.5 is expressed in cardiac progenitors and CMs and is required for cardiac development; however, Nkx2.5-deficient mice exhibited linear heart tube formation, suggesting that Nkx2.5 may not be required for cardiogenesis^27^. Blastocyst complementation in Nkx2.5-deficient mice consistently failed to generate hearts fully derived from exogenous PSCs^28^. In contrast, Mesp1/2-DKO mice exhibited mesodermal defects and lacked heart formation^18^. Here, we found that in Mesp1/2-DKO mice, CMs and non-CMs were virtually exclusively generated from PSCs via blastocyst complementation. Lineage tracing has revealed that most cardiovascular cells originate from the Mesp1^+^ cardiac mesoderm; nonetheless, a subset of cells, such as cardiac nerves and vascular SMCs, may be derived from neural crest cells^29^. Through blastocyst complementation, a combination of Mesp1/2 deletion and neural crest ablation may generate hearts derived exclusively from exogenous PSCs.

Interspecies blastocyst complementation using rat PSCs can generate a heart composed of rat-derived cells in Mesp1/2-DKO mouse. However, its efficiency is lower than that of the xenogenic pancreas and kidneys^6,7,9^. Higher chimerism may be necessary to generate the heart, the first organ to form during development. Using high-quality PSCs and establishing a permissive niche may facilitate the production of xenogenic hearts. Further studies are required; nevertheless, our findings reveal that the method has the potential to generate a rejection-free heart.

## Methods

### Animals

The Tsukuba University Ethics Committee for Animal Experiments approved this study. Mesp1/2 heterozygous KO mice (Mesp1^+/−^/Mesp2^+/−^, Mesp^+/−^) were provided by RIKEN BRC through the National BioResource Project of MEXT/AMED, Japan^18^. Intercrossing heterozygous Mesp^+/−^ mice yielded homozygous Mesp1/2-DKO (Mesp1^−/−^/Mesp2^−/−^, Mesp^−/−^) mice. Mesp1-Cre/R26-tdTomato mice were obtained by crossing Mesp1-Cre and tdTomato reporter mice^30,31^. Transgenic mice overexpressing GFP under the control of the CAG promoter into ROSA26 (R26-GRR) were described previously^32^. ICR (Crl: CD1(ICR)) mice were purchased from The Jackson Laboratory Japan (Yokohama, Japan).-Sprague Dawley (SD) Tg (CAG-EGFP) rats were purchased from Japan SLC, Inc. (Shizuoka, Japan). No immune deficiencies or other health problems were observed in the transgenic animals, and all animals were experimentally and drug-naïve before use. All animals were group-housed and bred in a dedicated husbandry SPF facility with 14 h/10 h light-dark cycles, *ad libitum* food, and water access, and were checked daily. To maintain their SPF grade, their health statuses were routinely checked. Under the same conditions, animals subjected to surgical procedures were transferred to a satellite-specific pathogen-free (SPF) facility.

### Rodent ESCs

GRR and SGE2 mESCs (GFP-expressing mESCs) have previously been described^6,32^. Undifferentiated GRR and SGE2 mESCs were maintained on mitomycin-c-treated mouse embryonic fibroblasts (MEFs) derived from ICR. The culture medium for GRR consisted of Dulbecco’s modified Eagle’s medium (DMEM; 11995, Thermo Fisher Scientific) supplemented with 10% KnockOut serum replacement (KSR; 10828, Thermo Fisher Scientific), 1% non-essential amino acids (1681049, MP Biomedicals), 0.1 mM 2-mercaptoethanol (M3148 Merck), and 1000 U/mL LIF (ESG1107, Merck), 3 μM CHIR99021 (04-0004-02, Reprocell), and 1 μM PD0325901 (04-0006-02, Reprocell), following standard procedures. The culture medium for SGE2 consisted of NDiff 227 (Y40002, Takara) supplemented with 1000 U/mL LIF (ESG1107, Merck), 3 μM CHIR99021 (04-0004-02, Reprocell), and 1 μM PD0325901 (04-0006-02, Reprocell) following standard procedures. Tetraploid complementation was used to validate all ESC lines^33^.

Rat ES cells (rESCs) were established using a previously described method^32,34^. The 3.5 dpc embryos were collected from pregnant rats and cultured for 1 d in KSOM (ARK Resource, Kumamoto, Japan). Blastocysts were placed on a feeder layer after the zona pellucida was removed with an acidic Tyrode solution (T1788, Sigma). Following trypsinization, the proliferating inner cell mass (ICM) was collected using a glass pipette and seeded on a new feeder layer on the fourth or fifth day. Furthermore, the established ESCs were collected on the fourth or fifth day. Only early passage rESCs were used for injection. The culture medium for rESCs consisted of NDiff 227 (Y40002, Takara) supplemented with 1000 U/mL LIF (ESG1107, Merck), 3 μM CHIR99021 (04-0004-02, Reprocell), and 1 μM PD0325901 (04-0006-02, Reprocell) following standard procedures.

### Blastocyst complementation using GFP-blastomeres

Crossing Mesp1/2 heterozygous KO (Mesp^+/−^) mice yielded Mesp^−/−^ embryos. Chimeric embryos were generated by aggregating eight-cell embryos from GRR mice with four-cell embryos from mutant mice. Briefly, four- and eight-cell stage embryos were obtained from *in vitro* fertilized (IVF) Mesp^+/−^ and GRR mice, respectively. Mutant chimeras were generated by combining one embryo (Mesp-/-, Mesp+/−, or Mesp+/+) with 1-2 GRR blastomeres. KSOM medium (MR-121-D; Merck) was used to culture the chimeric embryos. Twenty successfully aggregated chimeric blastocysts were implanted surgically into the uterine horns of pseudopregnant ICR females (2.5 dpc) and allowed to grow to the desired stage.

### Blastocyst complementation with ES cells

Embryo freezing was performed for 72 h following IVF and used for blastocyst complementation with ESCs. Host embryos were prepared by thawing frozen embryos and incubating them for 24 h in an M16 medium (M7292, Sigma-Aldrich). Further, 3 d before embryo manipulation, frozen ESCs were thawed and placed in a feeder layer. Proliferating ESCs were dissociated shortly before injecting with 0.25% trypsin. Under a microscope, a piezo-driven micromanipulator (PMM-150FU, PrimeTech) was used to perforate the zona pellucida and trophectoderm under a microscope, and 10–20 ESCs were injected into the blastocyst cavities near the inner cell mass using the injection method. Manipulated chimeric embryos were implanted at the blastocyst stage into the uterine horns of pseudopregnant ICR female mice (2.5 dpc).

### Mouse fibroblasts for genotyping

MEFs and tail-tip fibroblasts (TTFs) were isolated as previously described^35–37^. Briefly, embryos were extracted from pregnant mice and washed with phosphate-buffered saline (PBS). Each of the tissue of the embryos was washed in fresh PBS, chopped using a pair of scissors, transferred to a 0.25% trypsin/ethylenediaminetetraacetic acid (EDTA) solution (25200-072, Gibco), and incubated for 5 min at 37°C. Following trypsinization, twice as much MEF media (DMEM containing 20% fetal bovine serum (FBS) per embryo) was added and pipetted several times to facilitate tissue dissociation. The supernatant was transferred to new tubes, and cells were collected through centrifugation and they were resuspended in DMEM/20% FBS (CCP-FBS-BR-500; COSMO Bio) for culturing at 37°C in 5% CO_2_. A 1 cm tail of a 4-week-old mouse was minced into pieces < 3 mm. The minced tail was transferred to an enzyme digestion solution containing 1 mg collagenase D (11088866001, Merck) in 0.4 mL PBS and incubated at 37°C for 90 min. The digested tail tissue was transferred to a 70-μm cell strainer placed in a 10 cm cell culture dish with 10 ml of TTF medium (Iscove’s modified Dulbecco’s medium (IMDM; 12440053, Gibco) with 20% FBS) and vigorously pulverized with a syringe plunger. The supernatant was transferred to new tubes, and cells were collected by centrifugation and resuspended in a TTF medium for culturing at 37°C in 5% CO_2_.

### Genotyping of chimeric mice

The genotypes of the chimeric mice were confirmed using genomic DNA extracted from GFP-negative fibroblasts obtained using FACS (MoFlo XDP, Beckman Coulter). MEFs were isolated from the entire fetus and head tissues on embryonic days (E) 9.0~9.5 and after E10.5, respectively. After four weeks, adult TTFs were extracted from the tail. Detection of Mesp^−/−^ mutant alleles by genomic PCR was performed using four primer sets to amplify WT (439 and 829 bp PCR products) and KO (1.8 kbp PCR product): F1, 5’-GCCACCTGGCCCTTGCTACTCC-3,’ F2 5’-GAGCTGCACCTGGCCTGCAGTG-3’, R1 5’-CTGAGTTCATTGACGTGGACCC-3’, and R2 5’-GCTTTGGTCTTGGGCTTCCAGTG-3’.

### Fluorescence-activated cell sorting

MoFlo XDP was used for analyses and cell sorting. To exclude GFP-silenced immune cells, fibroblasts were stained with an APC-conjugated anti-CD45 antibody (103112, Biolegend, 1:100). FlowJo software (Becton, Dickinson and Company) was used to evaluate the data collected.

### Immunohistochemistry

Embryos were fixed in 4% paraformaldehyde (PFA) overnight at 4°C, then in 30% sucrose overnight at 4°C, before being embedded in OCT compound and frozen in liquid nitrogen. The hearts, intestines, kidneys, livers, and lungs of adult mice were fixed in 4% PFA overnight and embedded in OCT compound for freezing in liquid nitrogen. Cryostat microtome (CM3050S, Leica) was used to cut these samples into 7 µm pieces. Sections were stained with primary antibodies against α-actinin (A7811, Sigma, 1:800), α-SMA (A2547, Sigma, 1:400), CD31 (ab28364, Abcam, 1:10), GFP (598, MBL, 1:500, and D153-3, MBL, 1:200), and Vimentin (GP53, Progen, 1:100, and ab45939, Abcam, 1:100) followed by secondary antibodies conjugated with Alexa 488 or 546 and DAPI (D1306, Invitrogen, 1:100). Images were captured using an all-in-one fluorescence microscope (BZX810, Keyence) or confocal microscope (LSM800, Carl Zeiss). The number of GFP^+^ cells was automatically counted using a hybrid cell count application (BZ-H4C, KEYENCE) in the BZ-X Analyzer software (BZ-H4A, KEYENCE). In each section, the ratio of GFP^+^ cells was counted in 10 non-overlapping randomly selected fields. The measurements and calculations were performed blindly.

### qRT-PCR

Total RNA was extracted from whole embryos following a standard protocol. qRT-PCR was performed using a StepOnePlus Real-Time PCR system with TaqMan probes (Applied Biosystems) (Supplementary Table). The mRNA levels were normalized to those of *Gapdh*.

### Echocardiography

Transthoracic echocardiography (Vevo 2100, Visual Sonics) was used to assess cardiac function in 7- to 8-week-old mice. For echocardiographic examination, mice were anesthetized with low-dose isoflurane. Two-dimensional targeted M-mode traces were obtained at the papillary muscle level. The Teichholtz formula was used to compute the EF.

### Treadmill fatigue test

A treadmill fatigue test was performed after 3 d of acclimatization to treadmill exercise. As a warm up, the mice raced uphill on a 15° treadmill for 5 min at 10 m min^−1^, followed by an increase in speed by 2 m min^-1^ every 2 min. Fatigue was determined when the mice continuously remained on the shock grid for 10 s, measuring the running time to fatigue, and calculating the running distance.

### Electrocardiogram

Electrocardiograms of mice anesthetized with low-dose isoflurane were recorded using the electrode pads of the Vevo system and observed for a minimum of 30 s.

### Statistical analysis

Statistical parameters, including the number of samples (n), descriptive statistics (mean and standard error of the mean), and significance, are reported in the figures and legends. Generally, at least n = 3 was used for each time point in each experiment. The statistical significance of differences between groups was determined using Student’s t-test or one-way analysis of variance (ANOVA) with Tukey’s post-hoc test. Differences were considered significant at *P* values < 0.05. GraphPad Prism software was used for statistical analysis.

## Acknowledgments

We are grateful to Y. Saga for kindly providing the Mesp1/2-DKO mice, members of the Nakauchi lab for providing SGE2 mESC line and critical discussions on the manuscript. This work was supported by research grants from the Research Center Network for Realization of Regenerative Medicine (JP22bm1123012), the Japan Agency for Medical Research and Development, the Japan Foundation for Applied Enzymology, MSD Life Science Foundation, the Japan Society for the Promotion of Science (21K08072, 22H03065), and Takeda Science Foundation.

## Author contributions

Y.A. designed, performed, and analyzed all experiments and wrote the manuscript. T.S, and M.I. designed and analyzed all experiments and wrote the manuscript. R.F., Y.Y., T.A., K.N., and S.H. performed the in vivo experiments. Y.T., H.S., S.M., and S.T. performed embryo manipulation and establishment of PSCs. T.Y., H.M., K.H., and T.M. performed data analysis.

## Ethics declarations

### Competing interests declaration

The authors declare no competing financial interests.

### Data availability statement

All the data in this study are available from the corresponding author upon reasonable request.

### Code availability statement

This study did not use any original code.

## Extended Data Figure Legends

**Extended Data Fig. 1.**
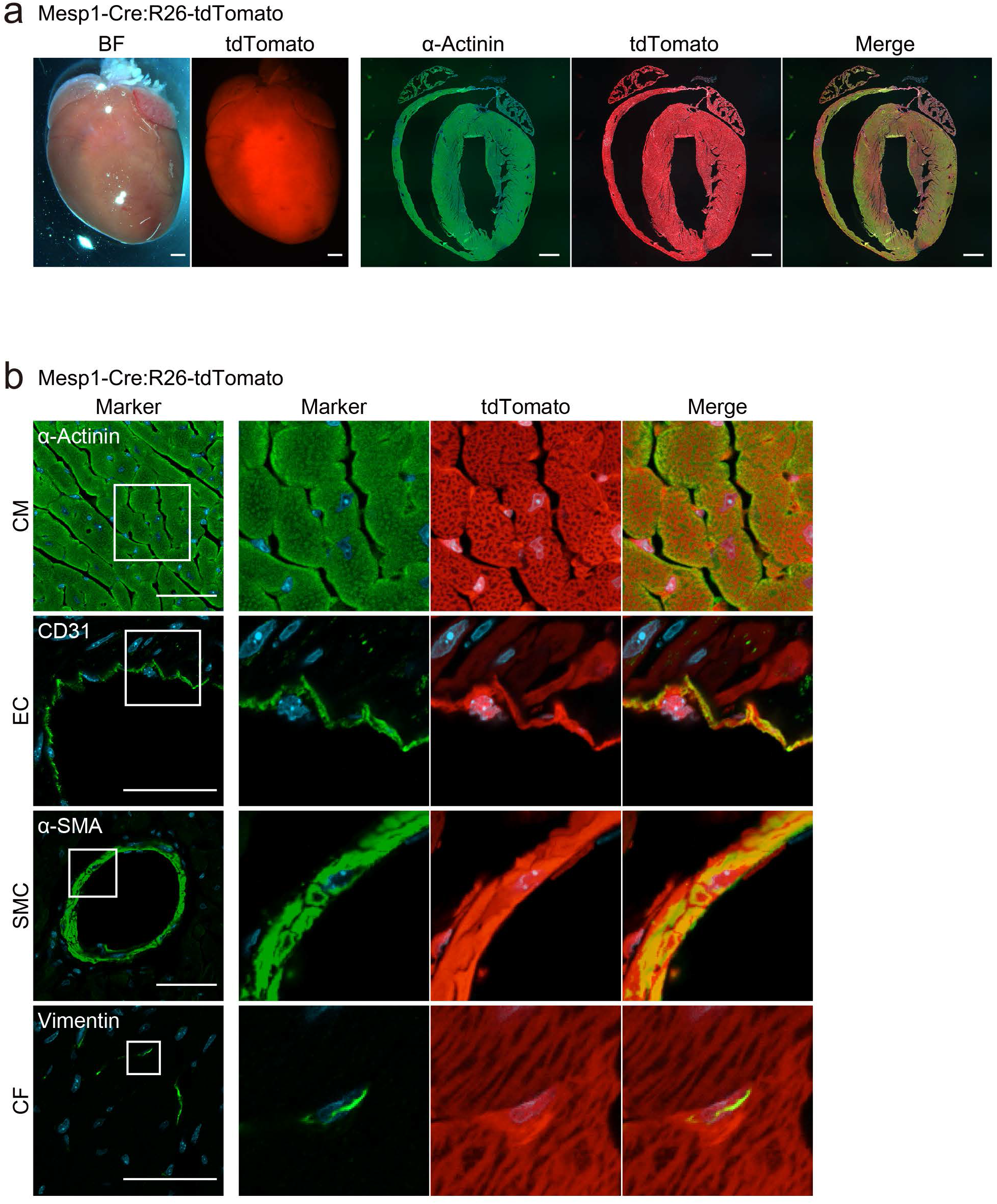
Lineage tracing of Mesp1+ cardiovascular progenitors. **a,** Whole-mount bright field and tomato fluorescence images and sections of Mesp1-Cre:R26-tdTomato mouse hearts. Heart sections were immunostained with α-actinin and DAPI. Note that there is a homogeneous tomato expression in the heart. Scale bars, 1 mm. **b,** Immunohistochemistry for the expression of cardiovascular cell markers and tdTomato in Mesp1-Cre:R26-tdTomato mouse hearts. α-actinin is a marker for cardiomyocytes (CM), CD31 for endothelial cells (EC), α-SMA for smooth muscle cells (SMC), and vimentin for cardiac fibroblasts (CF). High-magnification images are shown in the right panel. Scale bars, 100 μm.

**Extended Data Fig. 2.**
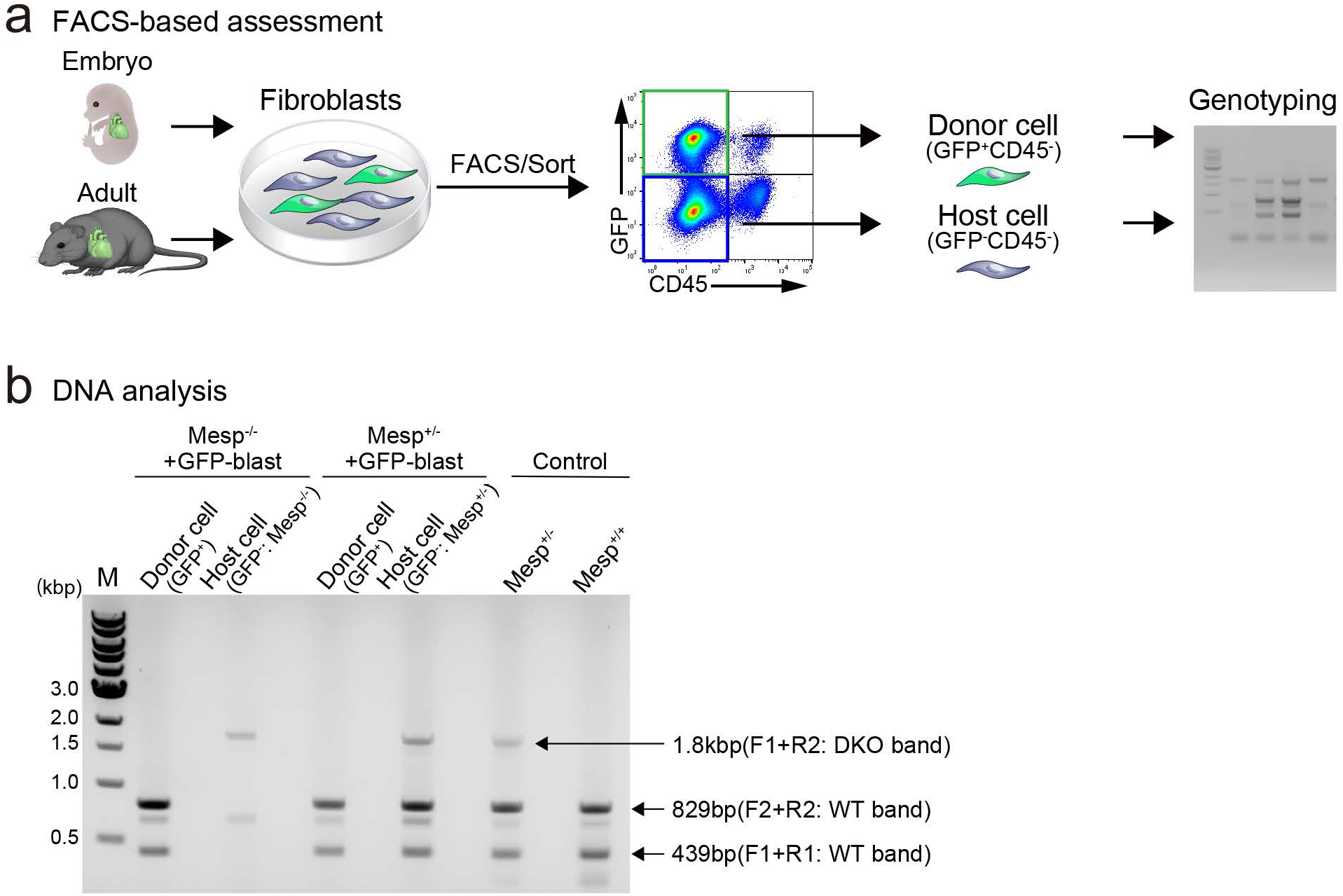
Genotyping strategy. **a,** Scheme of the FACS-based genotyping strategy. Donor-derived GFP^+^CD45^−^ fibroblasts and host-derived GFP^−^CD45^−^ fibroblasts were sorted and genotyped. **b,** Gel electrophoresis of PCR products from GFP^+^CD45 and GFP^−^CD45^−^ fibroblasts using the primers shown in (**Fig. 1a**). M, marker.

**Extended Data Fig. 3.**
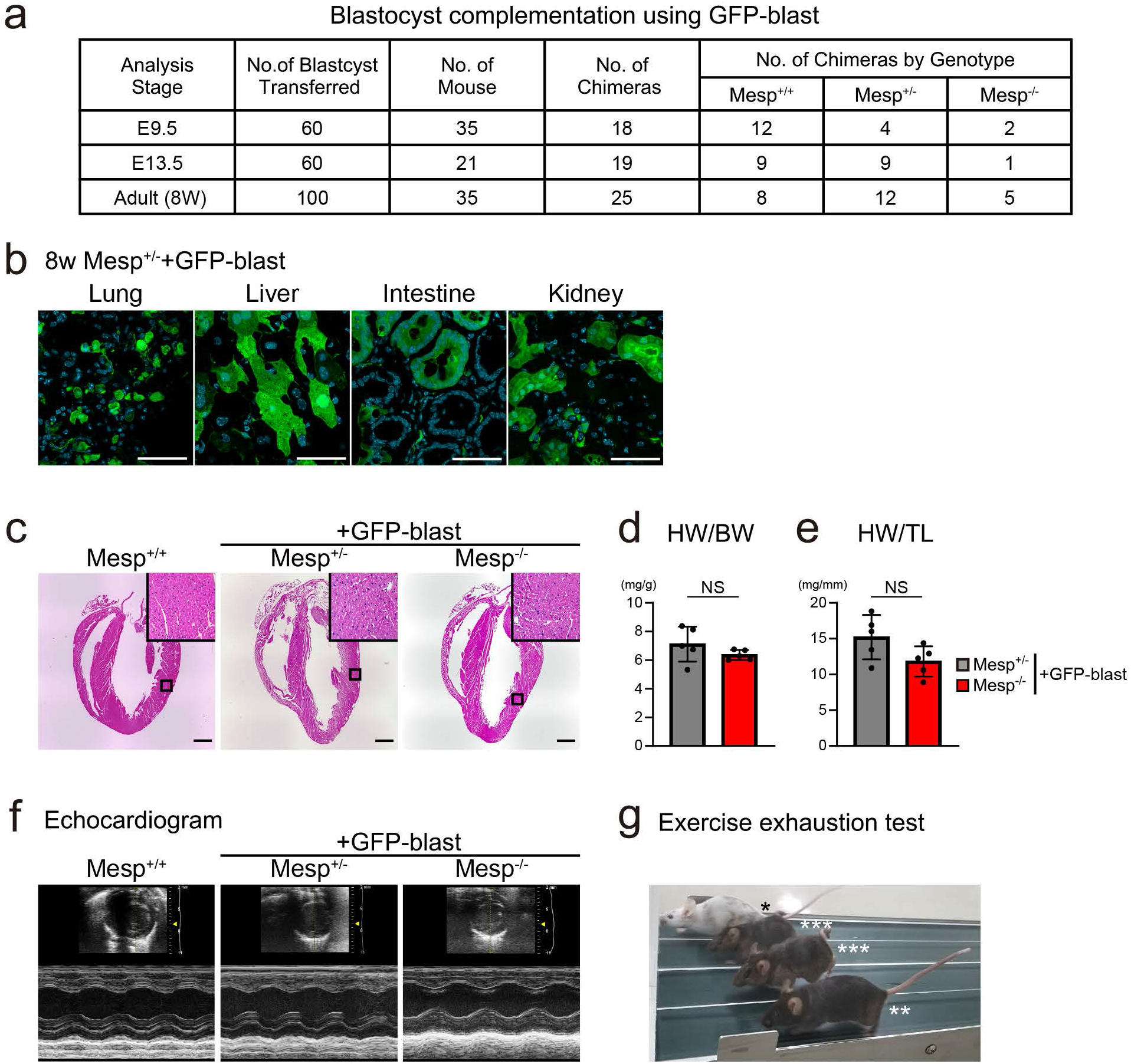
Blastocyst complementation using GFP-labeled blastomeres. **a,** Results of blastocyst complementation and genotyping. No., Number. **b,** Immunofluorescence images of GFP and DAPI in the lungs, liver, intestine, and kidneys of adult Mesp^+/−^ + GFP blast chimeras. Scale bars, 100 μm. **c,** Hematoxylin-eosin staining of 8-week-old mouse hearts (Mesp^+/+^, Mesp^+/−^ + GFP-blast, and Mesp^−/−^ + GFP-blast). High-magnification images are shown in the insets. Scale bars, 1 mm. **d, e,** Ratios of heart weight to body weight (HW/BW) (**d**) and heart weight to tibial length (HW/TL) (**e**). Data are presented as the mean ± SD of five mice (8-week-olds) per group. The Student’s t-test was used for statistical analysis. NS: statistically non-significant. **f,** Left ventricular M-mode echocardiographic images of Mesp^+/+^, Mesp^+/−^ + GFP-blast, and Mesp^−/−^ + GFP-blast mice. **g,** Image of treadmill fatigue test (*: Mesp^+/+^ + GFP-blast, **: Mesp^+/−^ + GFP-blast, ***: Mesp^−/−^ + GFP-blast).

**Extended Data Fig. 4.**
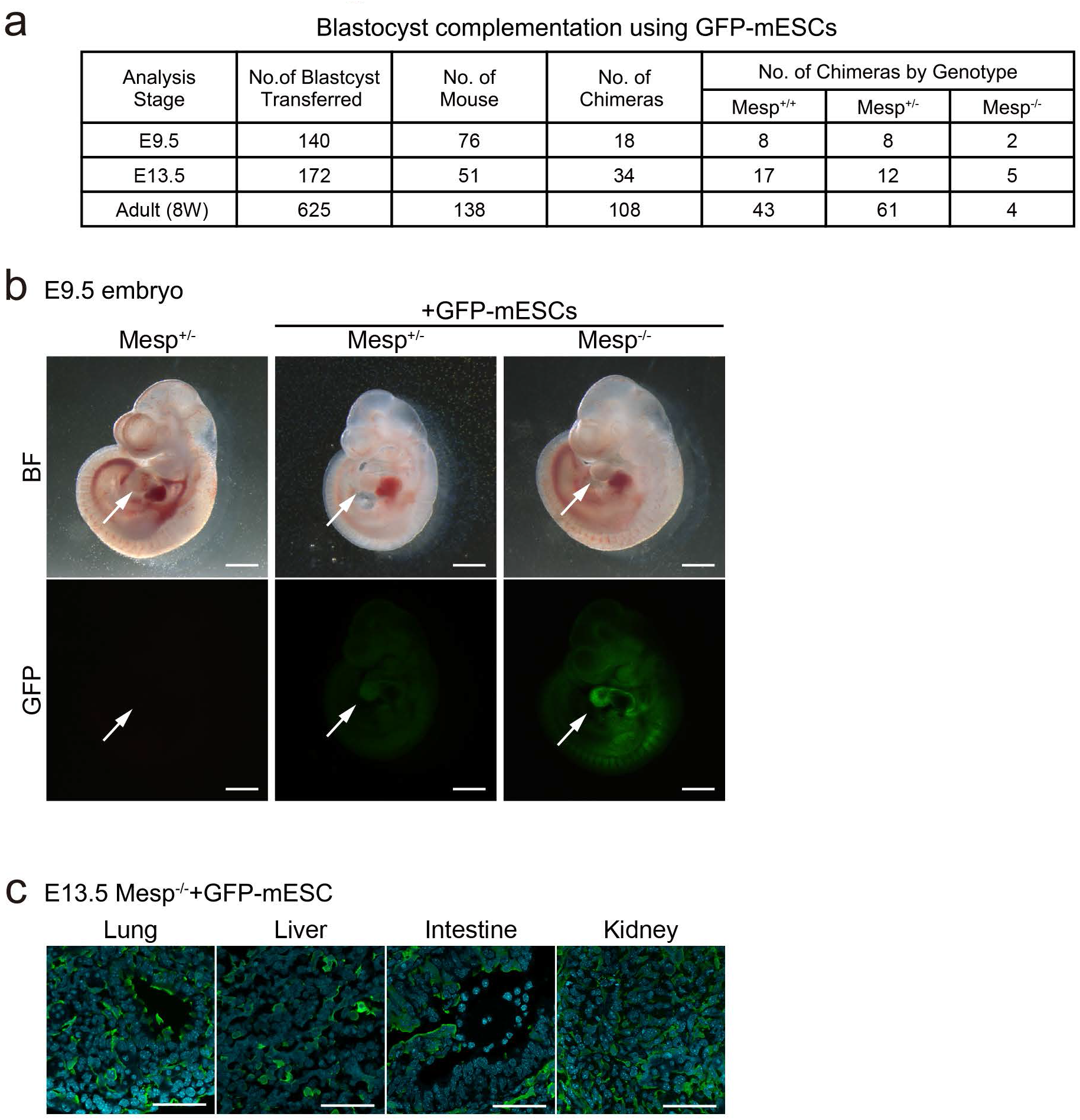
Blastocyst complementation using GFP-labeled mouse ES cells. **a,** Results of blastocyst complementation and genotyping. No., Number. **b,** Bright-field and GFP fluorescence images of E9.5 embryos (Mesp^+/−^, Mesp^+/−^ + GFP-mESCs, and Mesp^−/−^ + GFP-mESCs). Arrows indicate murine hearts. Scale bars, 500 μm. **c,** Immunofluorescence images of GFP and DAPI in the lungs, liver, intestine, and kidneys of E13.5 Mesp^−/−^ mice complemented with GFP-labeled mouse ESCs. Scale bars, 100 μm.

**Extended Data Fig. 5.**
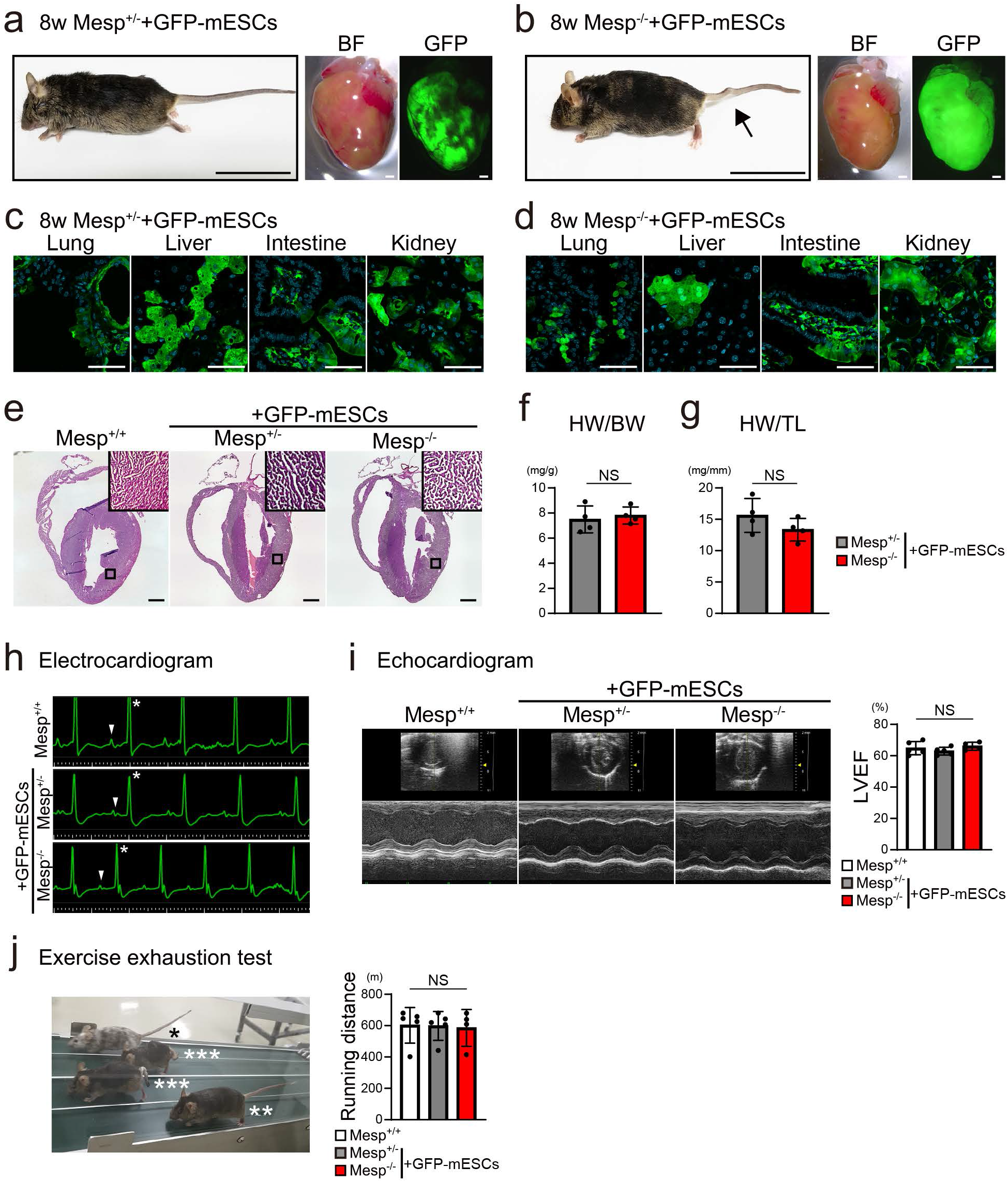
Generation of functional adult hearts by blastocyst complementation using mouse ES cells. **a, b,** Representative images of 8-week-old adult Mesp^+/−^ + GFP-mESCs (**a**) and Mesp^−/−^ + GFP-mESCs (**b**) mice and hearts. Whole-mount bright field and fluorescence images of the heart. The arrow indicates the kinked tail. Scale bars, 5 cm (mouse), and 1 mm (heart). **c, d,** Immunofluorescence images of GFP and DAPI in the lungs, livers, intestines, and kidneys of adult Mesp^+/−^ + GFP-mESCs (**c**) and Mesp^−/−^ + GFP-mESCs chimeras (**d**). Scale bars, 100 μm. **e,** Hematoxylin-eosin staining of hearts (Mesp^+/+^, Mesp^+/−^ + GFP-mESCs, and Mesp^−/−^ + GFP-mESCs). High-magnification images are shown in the insets. Scale bars, 1 mm. **f, g,** Ratios of heart weight to body weight (HW/BW) (**f**) and heart weight to tibial length (HW/TL) (**g**). Data are presented as the mean ± SD of four mice per group. The Student’s t-test was used for statistical analysis. NS: statistically non-significant. **h**, Electrocardiogram images of each mouse. Arrowheads, atrial P waves; * ventricular QRS waves. **I,** The left ventricular ejection fraction (LVEF) was quantified using echocardiography. Graph represents mean ± SD of n=5 (Mesp^+/+^), n=5 (Mesp^+/−^ + GFP-mESCs) and n=4 (Mesp^−/−^ + GFP-mESCs) mice per group. One-way ANOVA test with Tukey’s post-hoc test was used for statistical analysis. NS: statistically non-significant. **j,** Treadmill fatigue test and running distance. *, Mesp^+/+^; **, Mesp^+/−^ + GFP-mESCs; ***, Mesp^−/−^ + GFP-mESCs. Graph represents mean ± SD of n=5 (Mesp^+/+^), n=5 (Mesp^+/−^ + GFP-mESCs) and n=4 (Mesp^−/−^ + GFP-mESCs) mice per group. One-way ANOVA test with Tukey’s post-hoc test was used for statistical analysis. NS: statistically non-significant.

**Extended Data Fig. 6.**
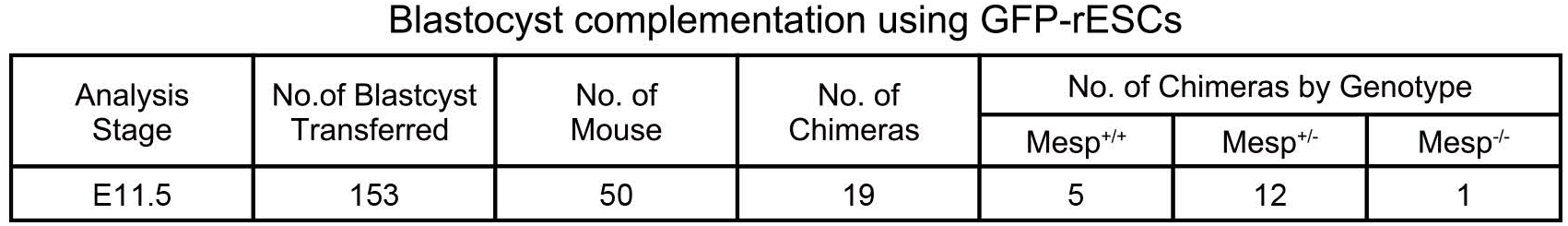
Interspecies blastocyst complementation using GFP-labeled rat ES cells. Results of blastocyst complementation and genotyping.

## Supplementary information

**Supplementary Table.**
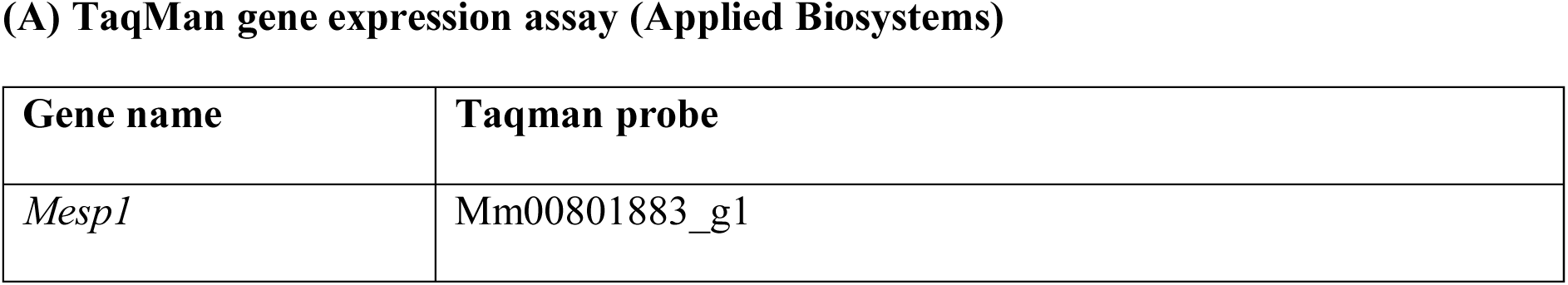
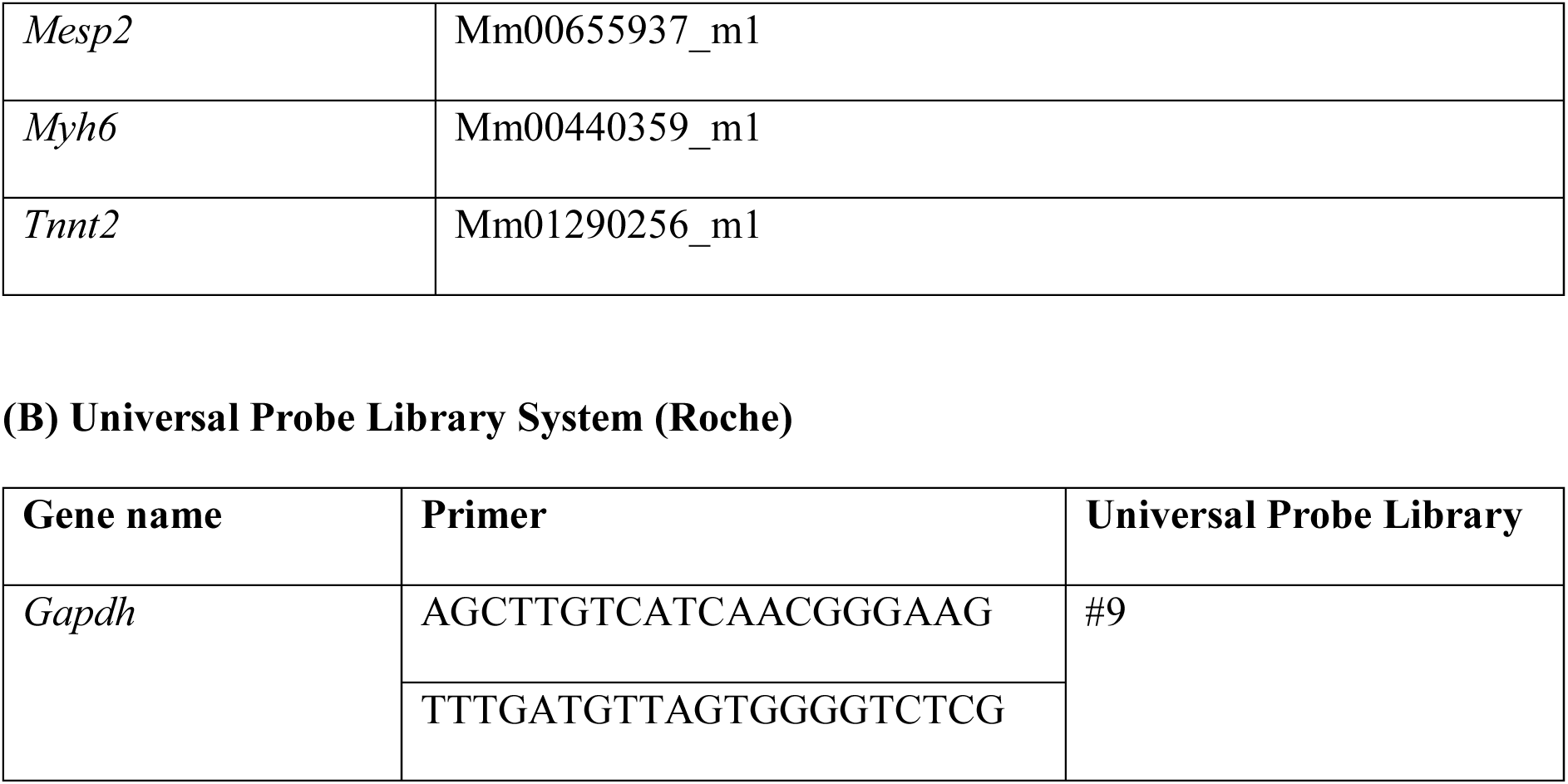
TaqMan probes (Applied Biosystems) and qRT-PCR primer sequences for Universal Probe Library (Roche), related to METHODS.

## Notes

### Competing Interest Statement

The authors have declared no competing interest.

